# Control of feeding by Piezo-mediated gut mechanosensation in *Drosophila*

**DOI:** 10.1101/2020.09.11.293712

**Authors:** Soohong Min, Yangkyun Oh, Pushpa Verma, David Van Vactor, Greg S.B. Suh, Stephen D. Liberles

## Abstract

Across animal species, meals are terminated after ingestion of large food volumes, yet underlying mechanosensory receptors have so far remained elusive. Here, we identify an essential role for *Drosophila* Piezo in volume-based control of meal size. We discover a rare population of fly neurons that express Piezo, innervate the anterior gut and crop (a food reservoir organ), and respond to tissue distension in a Piezo-dependent manner. Activating Piezo neurons decreases appetite, while *Piezo* knockout and Piezo neuron silencing cause gut bloating and increase both food consumption and body weight. These studies reveal that disrupting gut distension receptors changes feeding patterns, and identify a key role for *Drosophila* Piezo in internal organ mechanosensation.

## INTRODUCTION

Mechanosensory neurons detect a variety of environmental forces that we can touch or hear, as well as internal forces from organs and tissues that control physiological homeostasis (Abraira and Ginty, 2013; Ranade et al., 2015; Umans and Liberles, 2018). In many species, specialized mechanosensory neurons innervate the gastrointestinal tract and are activated by tissue distension associated with consuming a large meal (Williams et al., 2016; Zagorodnyuk et al., 2001). Gut mechanosensation may provide an evolutionarily conserved signal for meal termination, as gut distension inhibits feeding in many species and evokes the sensation of fullness in humans (Phillips and Powley, 1996; Rolls et al., 1998). However, how gut distension receptors contribute to long-term control of digestive physiology and behavior is unclear, as tools for selective pathway manipulation are lacking. Identifying neuronal mechanisms involved in detecting the volume of ingested food would provide basic insights into this fundamental mechanosensory process, and in humans, perhaps clinical targets for feeding and metabolic disorders.

Here, we investigated roles and mechanisms of food volume sensation in the fruit fly *Drosophila melanogaster*. Volumetric control of feeding was classically studied in a larger related insect, the blowfly, with relevant mechanosensory hotspots identified in the foregut and crop, an analog of the stomach (Dethier and Gelperin, 1967; Gelperin, 1967). In *Drosophila*, chemosensory neurons detect nutrients in the periphery and brain to control appetite, with some neurons positively reinforcing feeding during starvation conditions (Bjordal et al., 2014; Dus et al., 2015; Miyamoto et al., 2012). In contrast, the importance of gut mechanosensation in *Drosophila* feeding control and digestive physiology has not been similarly investigated; mechanosensory neurons of the gustatory system sense food texture and modulate ingestion (Sanchez-Alcaniz et al., 2017; Zhang et al., 2016), and other mechanosensory neurons in the posterior gut control defecation and food intake (Olds and Xu, 2014; Zhang et al., 2014). In contrast, food storage during a meal occurs primarily in the anterior gut (Lemaitre and Miguel-Aliaga, 2013; Stoffolano and Haselton, 2013). Enteric neurons of the hypocerebral ganglion innervate the fly crop, foregut, and anterior midgut, and lesioning of the recurrent nerve (which contains neurons of the hypocerebral ganglion) in *Drosophila* and blowfly increases feeding duration (Dethier and Gelperin, 1967; Gelperin, 1967; Pool et al., 2014). Together, these prior studies raise the possibility that a subpopulation of enteric neurons in *Drosophila* could be specialized to sense meal-associated gut distension.

## RESULTS

### Piezo-expressing enteric neurons innervate the gastrointestinal tract

To explore whether food volume sensation occurs in *Drosophila* and to investigate underlying mechanisms, we first asked whether neurons expressing various mechanosensory ion channels innervated the anterior gut. Several mechanosensitive ion channels have been reported in *Drosophila*, including TRP channels (Nompc, Nanchung, and Inactive), the degenerin/epithelial sodium channel Pickpocket (Ppk), transmembrane channel-like (Tmc) protein, and Piezo (Coste et al., 2012; Montell, 2005; Zhang et al., 2016; Zhong et al., 2010). We obtained Gal4 driver lines that mark neurons containing mechanoreceptor proteins or related family members, induced expression of membrane-tethered (CD8-GFP) or dendritically targeted (DenMark) fluorescent reporters, and visualized neuronal innervation of the anterior gut. We observed a small group of Piezo-expressing enteric neurons located in the hypocerebral ganglion (∼5-6 neurons per fly), and a dense network of Piezo fibers throughout the crop and anterior midgut (Figure 1A, 1B). Hypocerebral ganglion neurons were similarly labeled and anterior gut innervation similarly observed in three independent *Piezo-Gal4* driver lines (Figure S1A), but not in other Gal4 lines analyzed. We noted Nanchung expression in some epithelial cells of the crop duct, but not in crop-innervating neurons. The hypocerebral ganglion and adjacent corpora cardiaca together contain ∼35 neurons per fly based on Elav immunohistochemistry, and Piezo neurons therein were distinct from other neurons that expressed the fructose receptor Gr43a (∼5 neurons per fly) or the glucagon analog adipokinetic hormone (Akh, ∼20 neurons per fly) (Figure 1C, 1D, S1B). Piezo neurites formed a muscle-associated lattice in the gut and ascending axons contributed to the recurrent nerve (Figure 1E, 1F). *Drosophila* Piezo is expressed in many cell types (Kim et al., 2012), but its expression in gut-innervating sensory neurons was not reported previously. *Drosophila* Piezo was previously shown to confer mechanically activated currents when expressed in human cells, and to mediate mechanical nociception (Coste et al., 2012; Kim et al., 2012). Furthermore, vertebrate Piezo homologs play diverse mechanosensory roles, including in internal sensation of airway volume and blood pressure (Min et al., 2019; Nonomura et al., 2017; Zeng et al., 2018). We hypothesized that Drosophila enteric neurons which express Piezo and innervate the anterior gut might mediate volumetric control of appetite.

**Figure 1.**
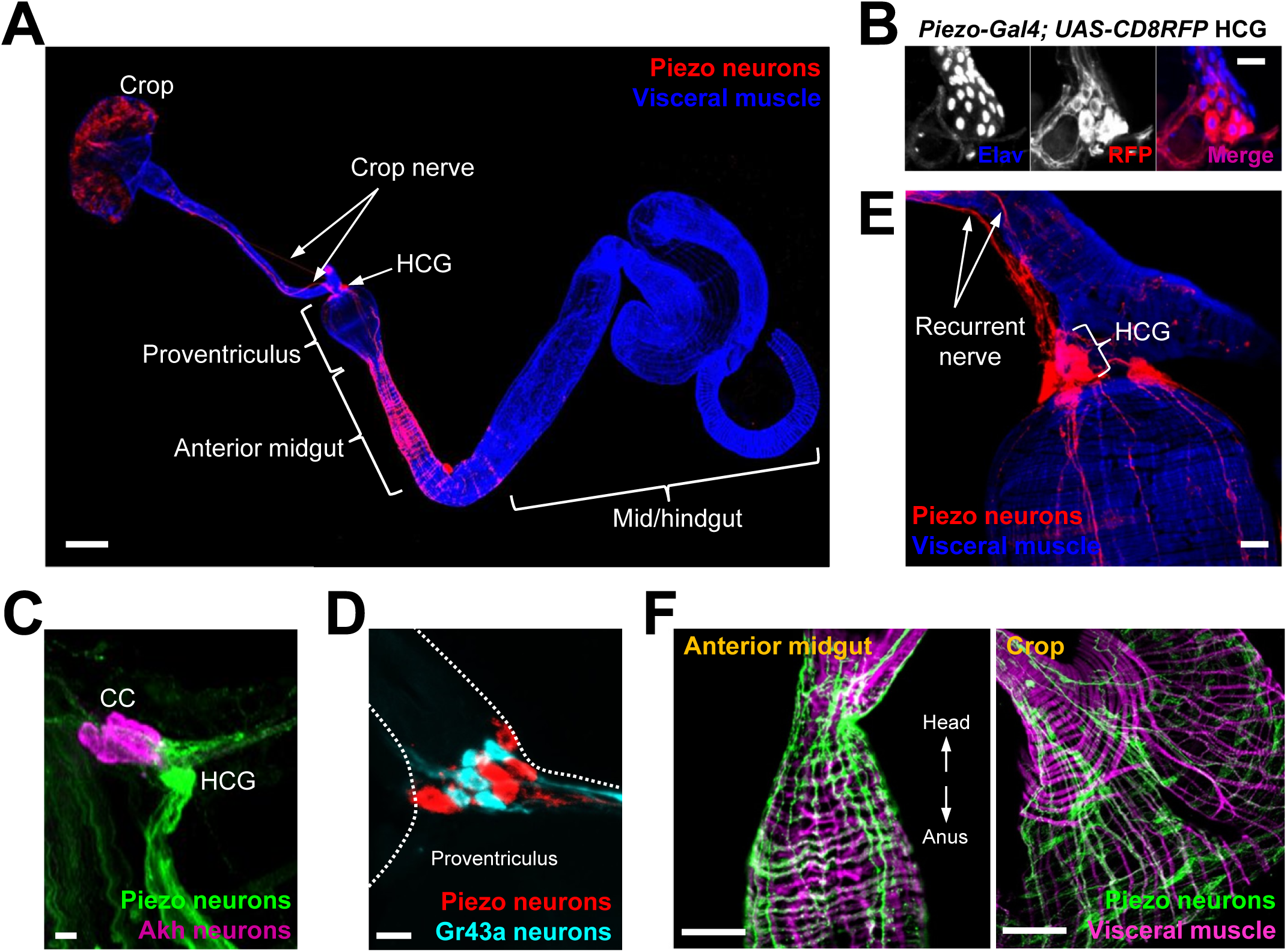
Piezo neurons innervate the gastrointestinal tract. (A) Wholemount image of the digestive tract from a *Piezo-Gal4 (59266); UAS-DenMark* fly visualized with immunofluorescence for DenMark (red, anti-RFP) and Phalloidin (blue) to label visceral muscle, HCG: hypocerebral ganglion, scale bar: 100 μm. (B) Immunofluorescence for RFP (red) and Elav (blue) in the HCG from a *Piezo-Gal4; UAS-CD8RFP* fly, scale bar: 10 μm. (C) Immunofluorescence for GFP (green) and Akh (magenta) in the corpora cardiaca (CC) and HCG from a *Piezo-Gal4; UAS-CD8GFP* fly, scale bar: 10 μm. (D) Native GFP and RFP fluorescence from the HCG of a *Piezo-Gal4; UAS-CD8RFP; Gr43a-LexA; LexAop-CD8GFP* fly, scale bar: 10 μm. (E) Image of the recurrent nerve (arrows) labeled by native RFP fluorescence in a *Piezo-Gal4; UAS-CD8-RFP* fly and Phalloidin immunofluorescence (blue), scale bar: 10 μm. (F) The anterior midgut (left) and crop (right) of a *Piezo-Gal4; UAS-DenMark* fly visualized by immunofluorescence for DenMark (green) and Phalloidin (magenta), scale bar: 50 μm.

### Piezo neurons control feeding behavior

To explore this model, we activated and silenced Piezo neurons using genetic approaches and monitored feeding behavior. We expressed temperature-sensitive Shibire (Shi^ts^) that blocks synaptic transmission at non-permissive temperatures (>32°C) in Piezo neurons using three independent *Piezo-Gal4* drivers (*Piezo>Shi*^*ts*^). *Piezo>Shi*^*ts*^ flies were reared at a permissive temperature (18°C) and later tested for physiological and behavioral changes at 32°C. To measure feeding behavior, flies were fasted for 24 hours, and then given brief access to food containing a dye for visualization and quantification of ingestion (Figure 2A). *Piezo>Shi*^*ts*^ flies from all three genotypes fed ravenously, and histological examination of the gastrointestinal tract showed gut bloating with increased crop size (Figure 2B). For comparison, genetic silencing of other gut-innervating neurons labeled in *GMR51F12-Gal4* flies did not impact appetite or cause crop distension. These findings indicate that disrupting Piezo neurons compromises gut volume homeostasis and associated control of feeding.

**Figure 2.**
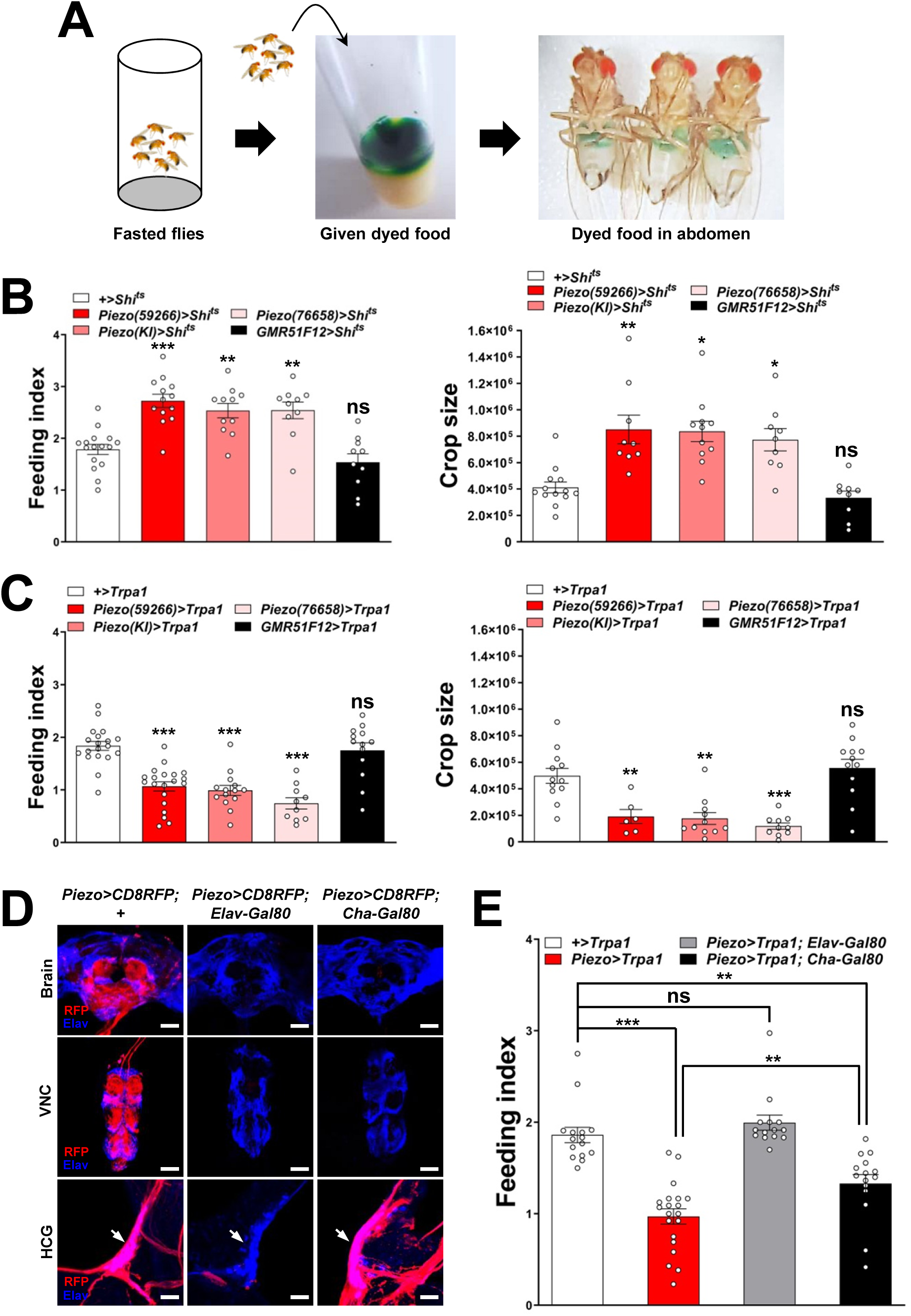
Piezo neurons control feeding behavior. (A) Depiction of the colorimetric feeding assay. (B) Fasted flies with *Shibire* alleles indicated were given brief access (30 min) to dye-labeled food at 32°C, and feeding indices and crop sizes were calculated, n (feeding index): 10-16 trials involving 120-192 flies, n (crop size): 9-13 flies, mean ± SEM, ***p<0.0005, **p<0.005, *p<0.05, ns: not significant by ANOVA Dunnett’s multiple comparison test. (C) Fasted flies with *Trpa1* alleles indicated were given brief access (30 min) to dye-labeled food at 30°C, and feeding indices and crop sizes were calculated, n (feeding index): 10-20 trials involving 120-240 flies, n (crop size): 6-12 flies, mean ± SEM, ***p<0.0005, **p<0.005, ns: not significant by ANOVA Dunnett’s multiple comparison test. (D) Native RFP fluorescence in brain (top), ventral nerve cord (VNC, middle) and hypocerebral ganglion (HCG, bottom) of *Piezo-Gal4*^*59266*^; *UAS-CD8RFP* flies with *Gal80* alleles indicated, scale bar: 100 μm (brain, VNC), 20 μm (HCG). (E) Fasted flies with *Trpa1* alleles indicated were given brief access (30 min) to dye-labeled food at 30°C, and feeding indices were calculated, n: 14-20 trials involving 168-240 flies, mean ± SEM, ***p<0.0005, **p<0.005, ns: not significant by ANOVA Dunnett’s multiple comparison test.

To test the effects of activating Piezo neurons on food consumption, we drove expression of the temperature-regulated ion channel Trpa1 in Piezo neurons using *Piezo-Gal4* lines (*Piezo>Trpa1*). Thermogenetic activation of Trpa1 in Piezo cells, achieved by transferring *Piezo>Trpa1* flies from 18°C to 30°C, suppressed food intake after a 24 hour fast, and also blocked meal-associated increases in crop volume, with similar results observed using three different *Piezo-Gal4* drivers (Figure 2C). Since many cell types express Piezo (Kim et al., 2012), we next used approaches for intersectional genetics involving Gal80, a dominant suppressor of Gal4-mediated gene induction to restrict Trpa1 expression to fewer cells. First, we drove Gal80 expression broadly in neurons using *Piezo>Trpa1*; *Elav-Gal80* flies, and observed restoration of normal feeding behavior, indicating the relevant Piezo expression site to be neurons (Figure 2D, 2E). Among neurons, *Piezo-Gal4* drove expression in various peripheral sensory neurons, the ventral nerve cord, brain, and hypocerebral neurons. Differential expression control could be partially achieved using a *Cha-Gal80* driver which silences Gal4-mediated expression in the ventral nerve cord and many central neurons, but not in gut-innervating hypocerebral neurons or a few cells of the proboscis, intestine, and brain (Figure 2D, S2A). Thermogenetic experiments in *Piezo>Trpa1*; *Cha-Gal80* flies also caused robust suppression of feeding behavior (Figure 2E). Intestinal cells are unlikely to contribute to feeding phenotypes in *Piezo>Trpa1*; *Cha-Gal80* flies based on experiments involving *Piezo>Trpa1*; *Elav-Gal80* flies; to provide additional evidence, thermogenetic activation of intestinal Piezo cells using *Escargot-Gal4; UAS-Trpa1* flies also had no effect on feeding (Figure S2B, S2C). In separate studies (Greg Suh, unpublished data), a role for central Piezo neurons in feeding regulation has also been observed, consistent with the significant differences we observe in feeding following thermogenetic activation experiments involving *Piezo-Gal4; UAS-Trpa1* and *Piezo-Gal4; UAS-Trpa1; Cha-Gal80* flies (Figure 2E). Additional studies are needed to isolate the contributions of hypocerebral ganglion Piezo neurons to feeding control.

### Piezo enteric neurons respond to crop-distending stimuli

Next, we investigated the response properties of Piezo-expressing enteric neurons. We analyzed neuronal activity using a transcriptional reporter system involving CaLexA through which sustained neural activity drives expression of GFP. CaLexA reporter was expressed in Piezo neurons using Gal4 drivers, along with an orthogonal activity-independent CD8-RFP reporter for normalization. For validation and determination of response kinetics, Trpa1-induced activation of Piezo neurons increased CaLexA reporter levels gradually, with maximal induction by 24 hours (Figure S3A). First, we asked whether hypocerebral Piezo neurons, and for comparison hypocerebral Gr43a neurons which function as peripheral sugar sensors, changed activity with feeding state (Figure 3A, 3B). For both neuron types, we observed that CaLexA-driven GFP expression was low after a fast or in flies fed *ad libitum*, but was strikingly elevated when flies engorged themselves on a sucrose diet (Figure 3A, 3B, S3B). Sucrose consumption could potentially stimulate both gut chemosensors and mechanosensors, as an increase in crop volume was observed compared with flies fed *ad libitum* (Figure S3C). We next asked whether activity changes in enteric neurons depended on the content of ingested material. We compared CaLexA-mediated GFP expression levels in flies fed for 24 hours with 1) sucrose, 2) sucralose, a sweetener that lacks caloric value and stimulates peripheral gustatory receptors but not internal Gr43a neurons, 3) water alone after a period of water deprivation, or 4) water alone *ad libitum*. Flies extensively consumed sucrose, sucralose, and water when water-deprived, resulting in acute increases in crop volume that were not observed in flies given only water *ad libitum* (Figure 3C). Enteric Gr43a neurons displayed elevated levels of CaLexA-mediated GFP expression after engorgement on sucrose, which is converted into fructose and glucose, but not sucralose or water, consistent with a role for these neurons in sensing nutritional carbohydrates (Miyamoto and Amrein, 2014). In contrast, enteric Piezo neurons were activated more generally by sucrose, sucralose, and deprivation-induced water ingestion, but not in controls given only water *ad libitum*, with responses correlated to the extent of gut distension. The observation that Piezo neurons were similarly activated by water-and sucrose-induced gut distension indicated a sensory mechanism that does not require chemosensation of particular nutrients. Together, these findings suggest a model of two segregated sensory pathways through the hypocerebral ganglion, with Gr43a neurons responding to sugars and Piezo neurons responding to anterior gut mechanosensation.

**Figure 3.**
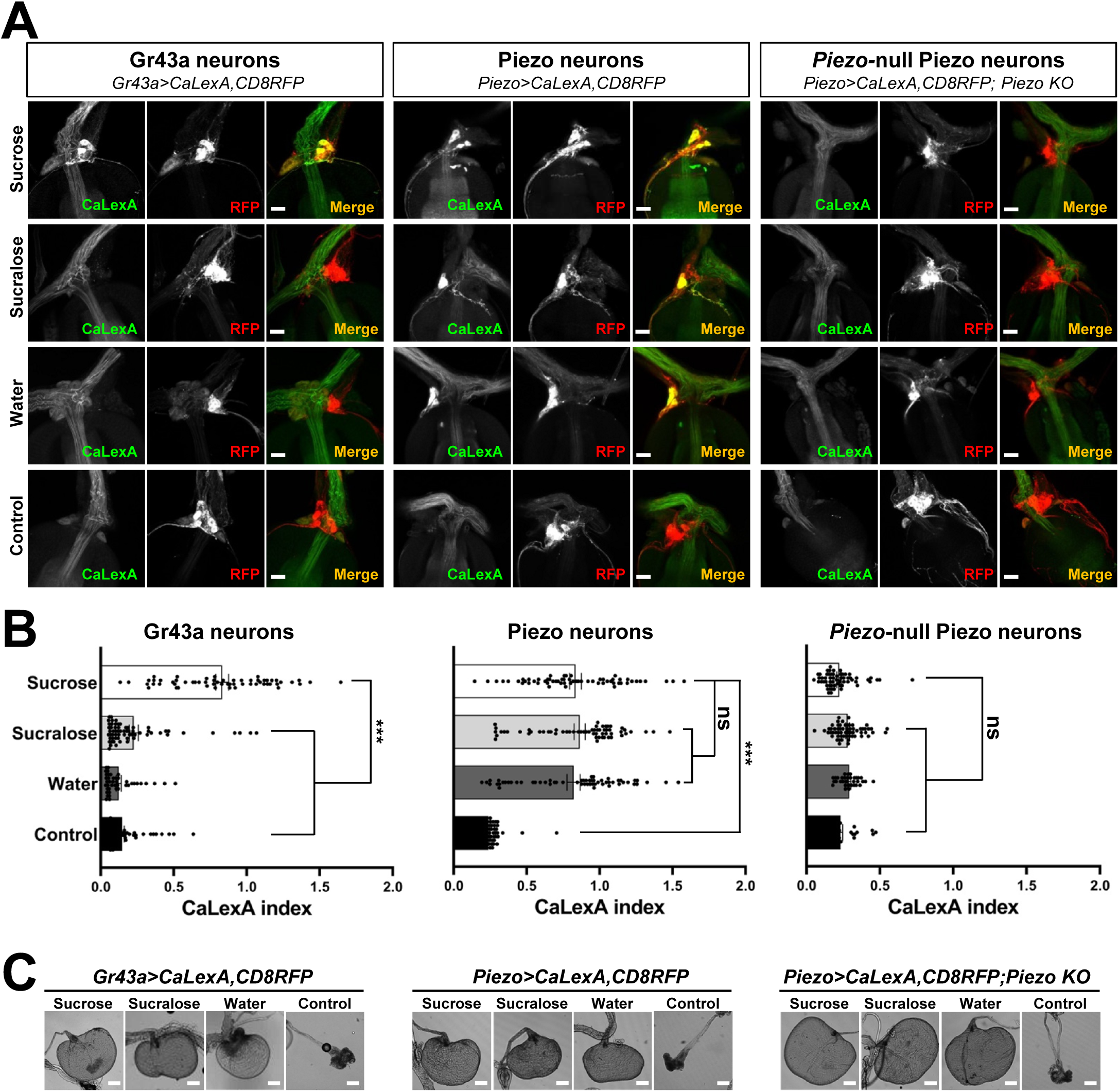
Piezo mediates enteric neuron responses to crop-distending stimuli. (A) Flies of genotypes indicated were provided solutions of 1) sucrose, 2) sucralose, 3) water alone after a period of water deprivation (water), or 4) water alone *ad libitum* for 24 hours (control). Representative images of native CaLexA-induced GFP reporter (green) and CD8RFP (red) fluorescence visualized in enteric Gr43a neurons (left), Piezo neurons (middle) or Piezo neurons lacking *Piezo* (right), scale bar: 10 μm. (B) Quantification of CaLexA-induced GFP fluorescence in individual RFP-expressing neurons from flies in A, n: 33-67 neurons from 5-15 flies, mean ± SEM, ***p<0.0001, ns: not significant by ANOVA Dunnett’s multiple comparison test. (C) Visualization of the crop from flies given stimuli indicated after 24 hours (sucrose, sucralose, control) or 15 minutes (water), scale bar: 100 μm.

### *Piezo* knockout alters enteric neuron responses and fly feeding behavior

Next, we asked whether the Piezo receptor mediates neuronal responses of hypocerebral neurons. We obtained Piezo knockout flies, and crossed them with flies harboring alleles enabling the CaLexA reporter system in Piezo neurons (using *Piezo-Gal4*^*59266*^ flies with the *Piezo-Gal4* transgene remote from the endogenous Piezo locus). Remarkably, hypocerebral ganglion neurons marked in *Piezo-Gal4* flies but lacking *Piezo* expression did not respond to engorgement by sucrose, sucralose or water, even though the crops of *Piezo* knockout flies were distended (Figure 3A, 3B). (As shown below, the extent of distension is actually more pronounced in *Piezo* knockout flies, yet CaLexA-mediated responses were not observed). A lack of neuronal responses in *Piezo* knockout flies is not due to gross deficits in the ability to produce reporter, as Trpa1-mediated activation of Piezo neurons in Piezo knockout flies was sufficient to induce a CaLexA-mediated response (Figure S3D). Furthermore, Piezo neurons displayed similar anatomical innervation of the anterior gut, suggesting that the deficit was not due to developmental miswiring (Figure S3E). Instead, enteric neurons of *Piezo* knockout flies seemingly fail to respond to crop-distending stimuli due to a mechanosensory defect.

Next, we asked whether *Piezo* knockout flies display changes in behavior or physiology. We measured feeding behavior in *Piezo* knockout flies, and for comparison, isogenic *w*^*1118*^ flies. For synchronization, flies were fasted for 18 hours and then given *ad libitum* access to dye-labeled food for 30 minutes. Remarkably, *Piezo* knockout flies increased food intake and had visually observable crop distension (Figure 4A, 4B, 4C). Moreover, *Piezo* knockout flies fed *ad libitum* on normal fly food for 5-7 days showed an increase in body weight compared to control flies (Figure 4D). Automated analysis of feeding patterns was performed involving an EXPRESSO platform, and *Piezo* knockout flies displayed an increase in food intake and feeding bout duration but a similar frequency of feeding bout initiation (Figure 4E). Abnormal gut distension and feeding behavior were rescued by exogenous expression of Piezo-GFP in *Piezo* knockout neurons driven by *Piezo-Gal4* (Figure 4F, S4A). Unlike Drop-dead knockout flies which have an enlarged crop due to defective food passage into the intestine (Peller et al., 2009), *Piezo* knockout flies have normal food transit, a normal lifespan, and increased defecation rates, presumably due to increased feeding (Figure S4B, S4C, S4D, S4E). Other than food-induced distension, the anatomy of the crop appeared normal in *Piezo* knockout flies, as visualized by histology of crop muscle, analysis of cell density, and volume measurements during starvation (Figure S4F, S4G, S4H). As mentioned above, knockout of *Piezo* does not impact the extent of gut innervation (Figure S3E); furthermore, thermogenetic Trpa1-mediated activation of Piezo neurons in *Piezo* knockout flies suppressed feeding behavior (Figure S4I), indicating that neural circuits downstream of enteric Piezo neurons were intact and remained capable of eliciting a behavioral response after *Piezo* knockout. Piezo also functions to guide the differentiation of gut enteroendocrine cells from mechanosensitive intestinal stem cells (ISCs) (He et al., 2018); however selectively restoring *Piezo* expression in ISCs using *Escargot-Gal4* (*Esg-Gal4*) that broadly marks Piezo-expressing ISCs did not rescue crop volume and feeding phenotypes (Figure S4J). We also note that while the crops of Piezo knockout flies are distended, the flies generally do not burst and do stop eating (although bursting does rarely occur), suggesting eventual engagement of a secondary satiety pathway, perhaps through nutrient sensors or posterior gut mechanoreceptors. Taken together, our data indicate a role for Piezo in sensing anterior gut distension, and that disrupting the function of Piezo neurons, or Piezo itself, causes substantial changes to gut physiology and feeding behavior.

**Figure 4:**
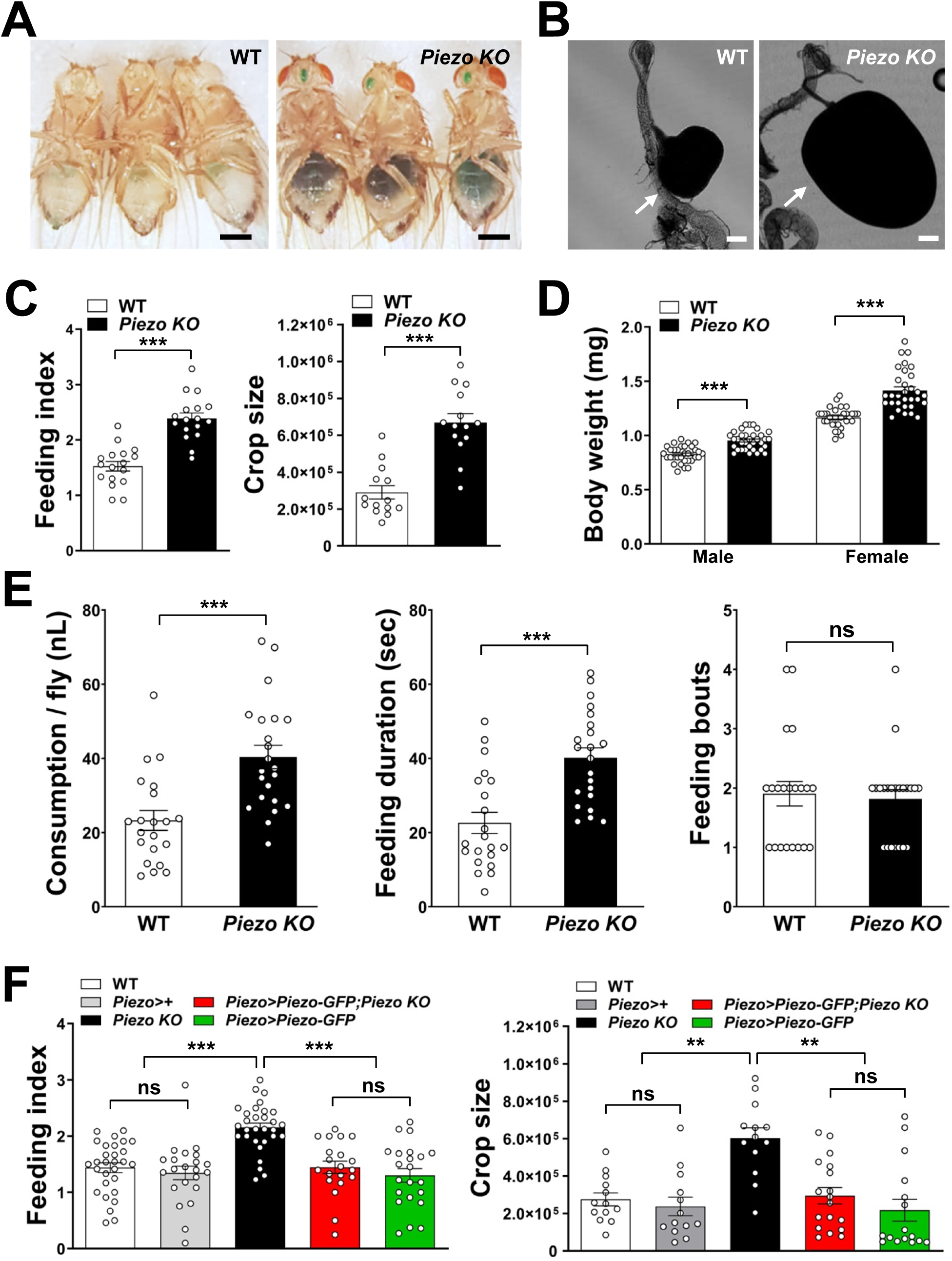
*Piezo* knockout alters fly feeding behavior. (A) Fasted wild type and *Piezo* knockout female flies were given brief access to dye-colored food and imaged, scale bar: 0.5 mm. (B) Representative images of the crop (arrow) in wild type and *Piezo* knockout flies, scale bar: 100 μm, (C) Calculated feeding indices (left) and crop sizes (right) from flies in A, n (feeding index: 17 trials involving 204 flies, n (crop size): 14 flies, mean ± SEM, ***p<0.0001 by unpaired t-test. (D) Body weights of wild type and *Piezo* knockout flies fed regular food *ad libitum*, n: 31-34 trials involving 93-102 flies, mean ± SEM. ***p<0.0001 by unpaired t-test. (E) Feeding parameters of fasted wild type and *Piezo* knockout male flies was analyzed using the EXPRESSO assay for 30 minutes after food introduction to determine overall food consumption, feeding duration per bout, and the number of bouts, n: 21-22 flies, mean ± SEM, ***p<0.0005, ns: not significant by unpaired t-test. (F) Calculated feeding indices (left) and crop sizes (right) from Piezo rescue and control flies indicated, n (feeding index): 20-30 trials involving 240-260 flies, n (crop size): 13-18 flies, mean ± SEM, ***p<0.0005, **p<0.005 by ANOVA Dunnett’s multiple comparison test, ns: not significant by unpaired t test.

Food-induced gut distension is thought to be an evolutionarily conserved signal for meal termination, yet underlying mechanisms and sensory receptors have long remained mysterious. Furthermore, whether food volume sensors are required for normal feeding control has remained unknown, as tools for selective loss-of-function were not available without knowing underlying sensory mechanisms. Here, we reveal a role for *Drosophila* Piezo in neurons that innervate the anterior gut and sense the size of a meal. Disrupting this pathway increases food consumption and body weight, and causes swelling of the gastrointestinal tract. These studies demonstrate that anterior gut mechanosensation contributes to the complex calculus that underlies the decision to eat, and provide a foundation for the comparative physiology and evolution of feeding control. Moreover, understanding related pathways in humans may enable new therapies for treating obesity and other food consumption disorders.

## ACKNOWLEDGMENTS

We thank Norbert Perrimon, Bryan Song, Dragana Rogulja for reagents and advice, Jinfei Ni for blinded analysis of behavior, Norbert Perrimon, Craig Montell, Julie Simpson, Hubert Amrein, and Bloomington Drosophila Stock Center for flies, and Hansine Heggeness and Exelixis facility at Harvard Medical School for fly food and stock maintenance. SM is supported by an American Heart Association postdoctoral fellowship (20POST35210914); PV and DVV are supported by NINDS award NS090994; GSBS is supported by NIH R01 grants (RO1DK116294, RO1DK106636) and Samsung Science and Technology Foundation (SSTF-BA-1802-11). SDL is an investigator of the Howard Hughes Medical Institute.

## AUTHOR CONTRIBUTIONS

SM, SDL, and GSBS designed experiments and analyzed data, SM and SDL wrote the manuscript, SM performed all experiments except YO performed EXPRESSO analyses and Akh co-staining, and PV and DVV assisted with fly husbandry.

## FIGURE LEGENDS

**Figure S1.**
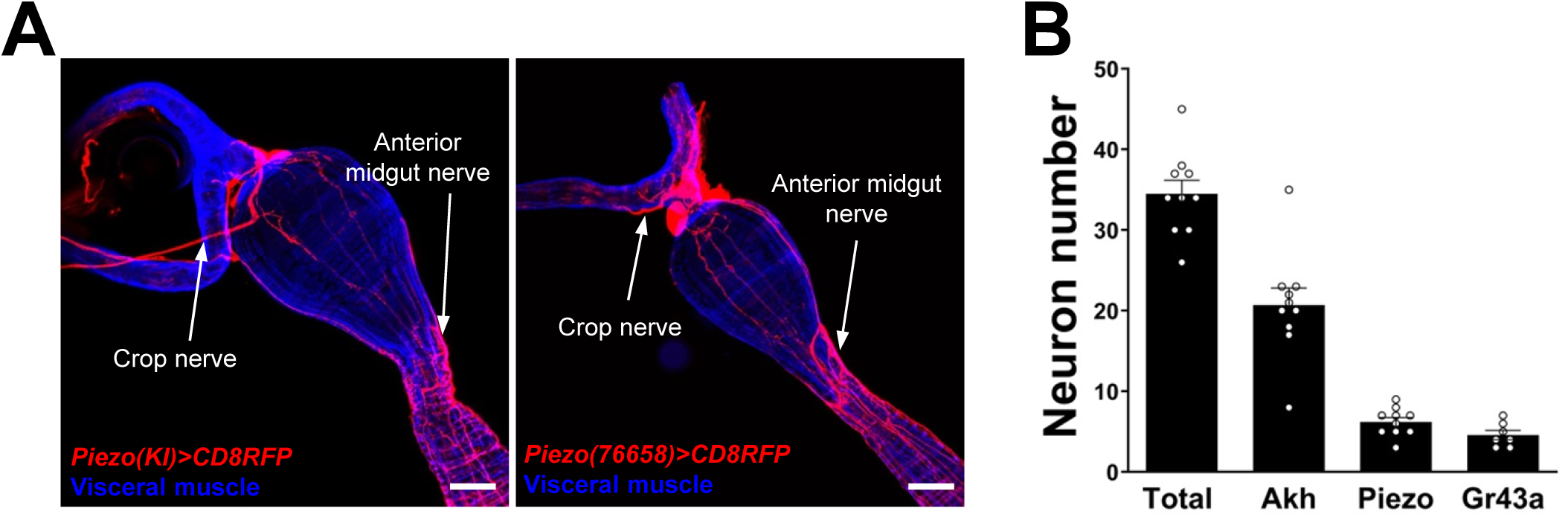
Innervation of the gastrointestinal tract by Piezo neurons. (A) Wholemount image of the digestive tract from two additional *Piezo-Gal4; UAS-CD8RFP* fly lines visualized with native RFP fluorescence (red) and Phalloidin immunofluorescence (blue) to label visceral muscle, HCG: hypocerebral ganglion, scale bar: 50 μm. (B) The average number of neurons in the hypocerebral ganglion and corpora cardiaca labeled per fly by immunofluorescence of Elav (total), Akh (AKH) and RFP (Piezo) in *Piezo-Gal4; UAS-CD8RFP* flies, and native GFP fluorescence (Gr43a) in *Gr43a-LexA; LexAop-CD8GFP* flies, n: 7-10 flies, mean ± SEM.

**Figure S2.**
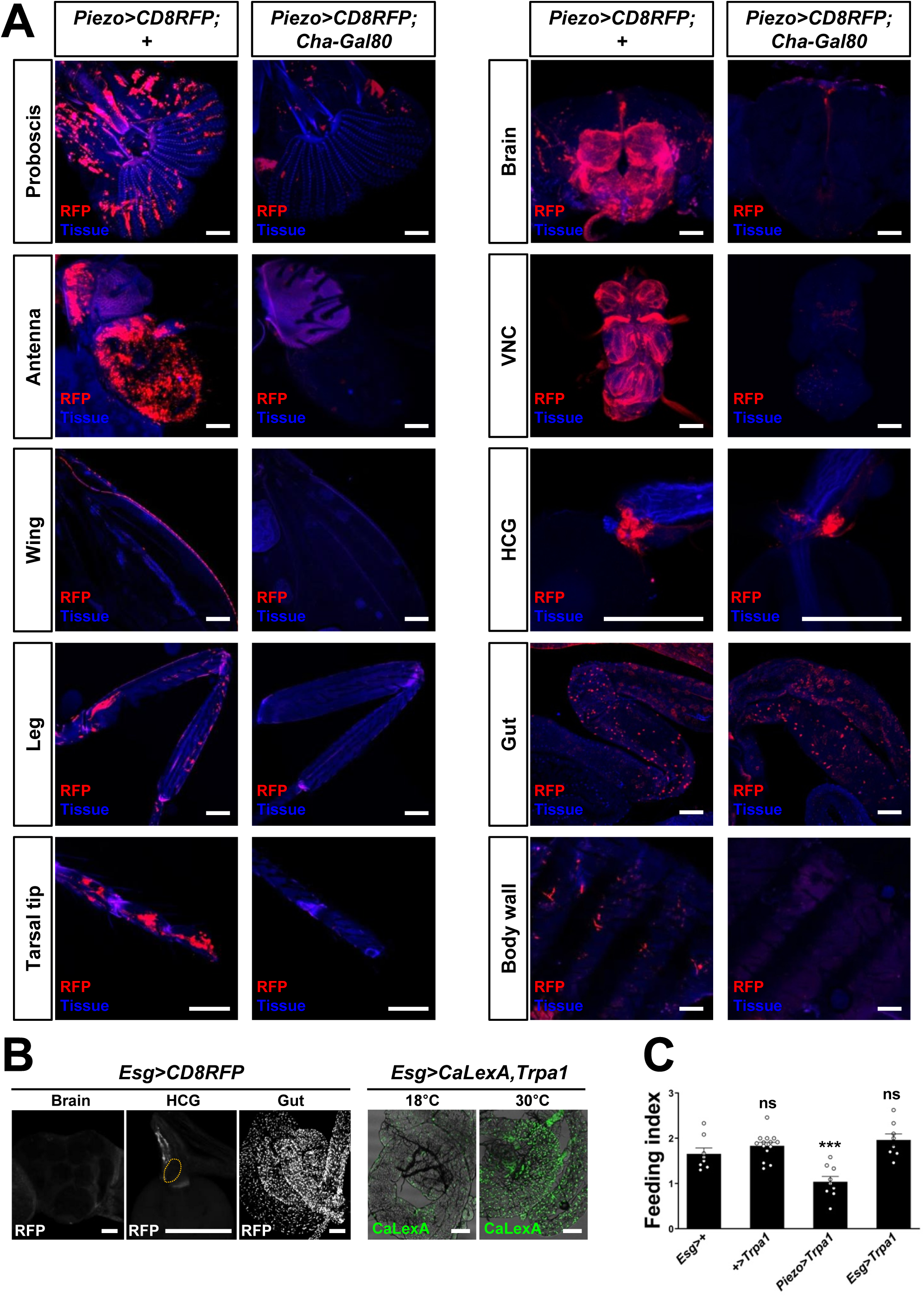
Visualizing and manipulating subtypes of Piezo neurons. (A) Native RFP fluorescence in tissues indicated from *Piezo-Gal4 (59266); UAS-CD8RFP* flies with or without *Cha*-*Gal80*, scale bars: 100 μm. (B) Wholemount images of native RFP fluorescence in the brain, proventriculus (orange outline: HCG) and midgut of *Esg-Gal4; UAS-CD8RFP* flies (left), and wholemount images of native GFP fluorescence in the midgut of *Esg-Gal4, UAS-CaLexA, UAS-Trpa1* flies at 18°C and 30°C (right). Scale bars: 100 µm. (C) Fasted flies with alleles indicated were given brief access (30 min) to dye-labeled food at 30°C, and feeding indices were calculated, n: 8-14 trials involving 96-168 flies mean ± SEM, ***p<0.0005, ns: not significant by ANOVA Dunnett’s multiple comparison test.

**Figure S3.**
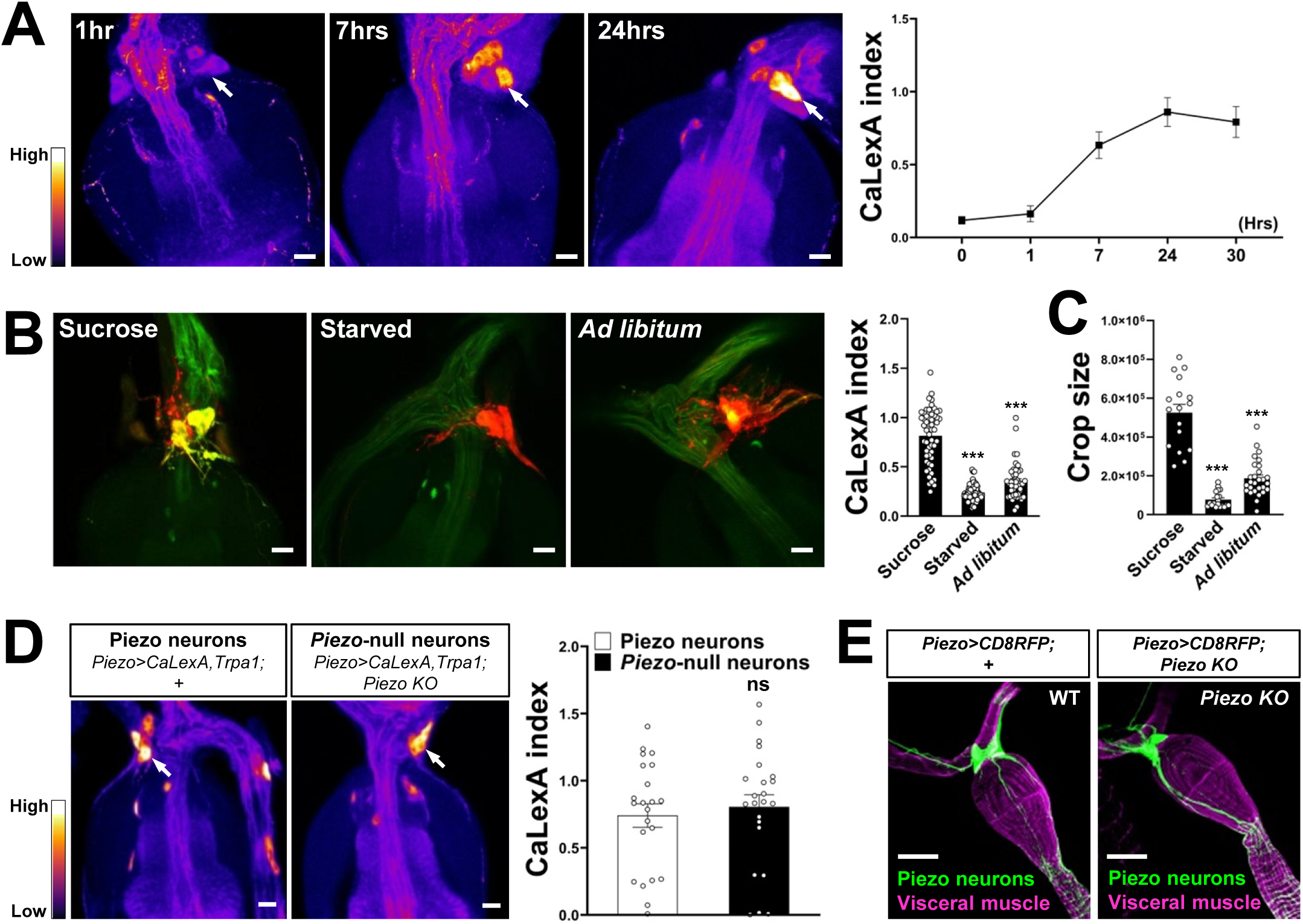
Responses and innervation patterns of Piezo neurons in wild type and Piezo knockout flies. (A) *Piezo-Gal4*; *UAS-CaLexA; UAS-Trpa1* flies were placed at 30°C for indicated time periods with *ad libitum* food. Representative pseudocolor images (left) and quantification (right) of native CaLexA-induced GFP reporter fluorescence in the hypocerebral ganglion. GFP fluorescence was visually transformed to a color map indicating fluorescence intensity, n: 10-18 flies, mean ± SEM, scale bar: 10 μm. (B) *Piezo-Gal4*; *UAS-CaLexA; UAS-CD8RFP* flies were provided for 24 hours with sucrose, water alone (starved) or regular food *ad libitum* (*Ad libitum*). Representative images (left) of native CaLexA-induced GFP reporter (green) and CD8RFP (red) fluorescence, and quantification (right) of CaLexA-induced GFP fluorescence in individual RFP-expressing neurons, n: 44-82 neurons from 8-15 flies, mean ± SEM, ***p<0.0001, ns: not significant by ANOVA Dunnett’s multiple comparison test, scale bar: 10 μm. (C) Crop sizes from wild type flies fed as indicated, n: 17-28 flies, mean ± SEM, ***p<0.0001 by ANOVA Dunnett’s multiple comparison test. (D) *Piezo-Gal4*; *UAS-CaLexA; UAS-Trpa1* (Piezo neurons) and *Piezo-Gal4*; *UAS-CaLexA; UAS-Trpa1; Piezo knockout* (*Piezo*-null neurons) flies were placed at 30°C with ad libitum food for 24 hours. Representative pseudocolor images (left) and quantification (right) of native CaLexA-induced GFP reporter fluorescence in the hypocerebral ganglion. GFP fluorescence was visually transformed to a color map indicating fluorescence intensity, n: 22-23 flies, mean ± SEM, ns: not significant by unpaired t-test, scale bar: 10 μm. (E) Wholemount images of the digestive tract from *Piezo-Gal4; UAS-CD8RFP* and *Piezo-Gal4; UAS-CD8RFP; Piezo knockout* flies visualized with immunofluorescence for RFP (green, anti-RFP) and Phalloidin (magenta) to label visceral muscle, scale bar: 50 μm.

**Figure S4.**
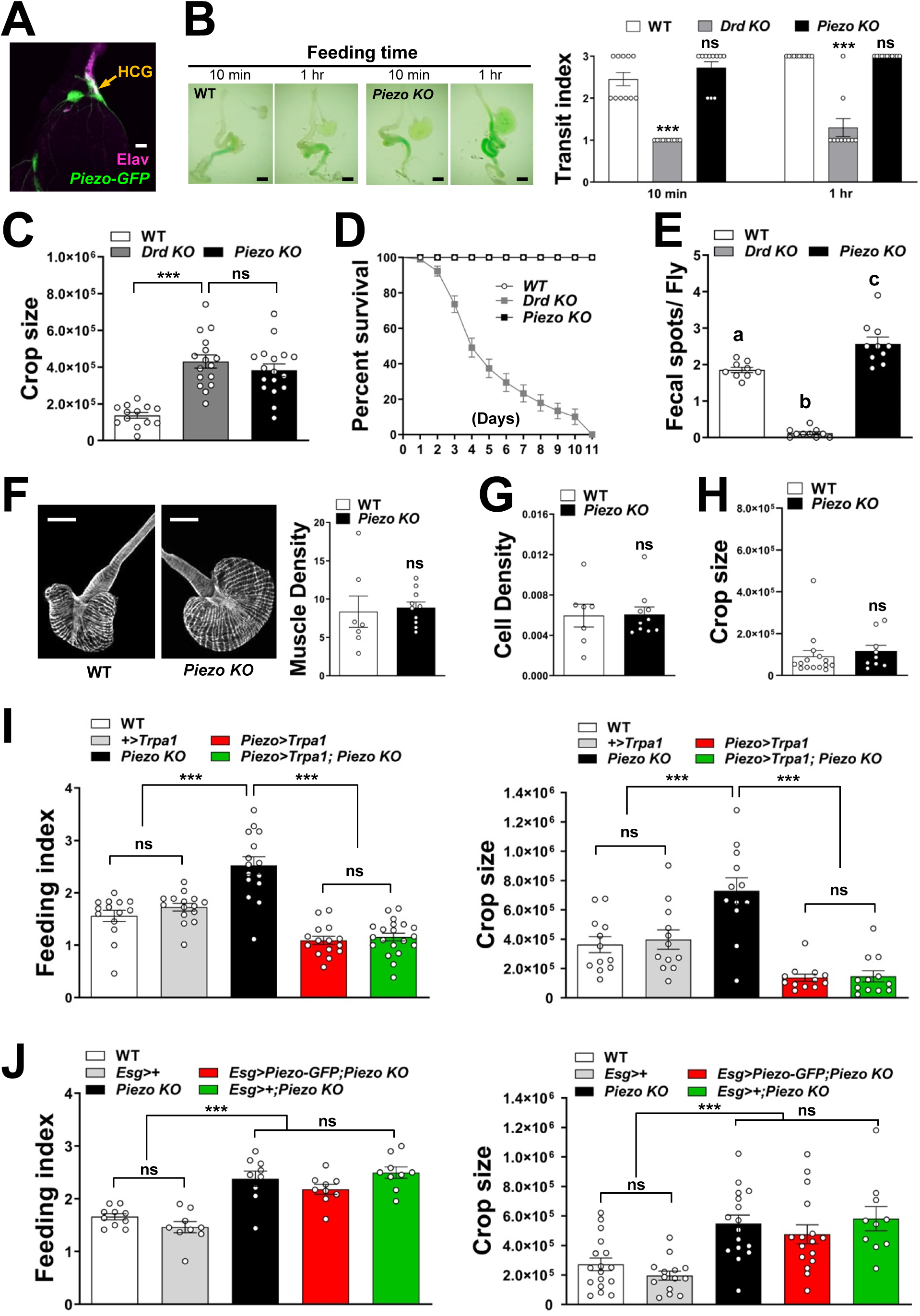
Physiological characterization of Piezo knockout flies. (A) Immunofluorescence for GFP (green) and Elav (magenta) in the hypocerebral ganglion (HCG) of Piezo rescue flies *[Piezo knockout; Piezo-Gal4 (59266); UAS-Piezo-GFP]*, scale bar: 20 μm. (B) Visualizing (left) and quantifying (right) intestinal transit of dye-colored food in wild type (WT), *Drop-dead* knockout (*Drd KO)* and *Piezo* knockout (*Piezo KO*) flies, n: 10-11 flies, mean ± SEM, scale bar: 200 μm. Measurements of (C) crop size, (D) survival, and (E) fecal spot deposition in wild type, *Piezo* knockout, and *Drop-dead* knockout flies, n for C: 13-17 flies, n for D: 7-8 trials involving 84-96 flies, n for E: 9-10 trials involving 90-100 flies, mean ± SEM, ***p<0.0001; ns: not significant by ANOVA Dunnett’s multiple comparison test; different alphabets indicate statistical difference. (F) Visualization (left) and quantification (right) of muscle fiber density by immunofluorescence for Phalloidin, n: 7-10 flies, mean ± SEM, scale bar: 100 μm. (G) Quantification of crop cell density based on nuclear staining (TO-PRO-3), n: 7-10 flies, mean ± SEM. (H) Measurement of crop size in fasted flies, n: 9-15 flies, mean ± SEM. (I) Fasted flies of genotypes indicated were given brief access (30 min) to dye-labeled food at 30°C, and feeding indices and crop sizes were calculated, n (feeding index): 14-20 trials involving 168-240 flies, n (crop size): 12 flies, mean ± SEM, ***p<0.0005, ns: not significant by ANOVA Dunnett’s multiple comparison test or unpaired t-test. (J) Fasted flies of genotypes indicated were given brief access (30 min) to dye-labeled food, and feeding indices and crop sizes were calculated, n (feeding index): 9-10 trials involving 108-120 flies, n (crop size): 10-17 flies, mean ± SEM, ***p<0.0005, ns: not significant by ANOVA Dunnett’s multiple comparison test.

## MATERIALS AND METHODS

### Flies

Fly stocks were maintained on a regular cornmeal agar diet (Harvard Exelixis facility) at 25°C, with mating and collection performed under CO_2_ anesthesia. For *Piezo* knock-out studies, *Piezo* knockout flies were isogenized by outcrossing five times into a wild type *w*^*1118*^ isogenic background. We obtained *Piezo* knockout, knock-in (KI) *Piezo-Gal4* and *UAS-Piezo-GFP* flies (Norbert Perrimon), *Tmc-Gal4* (Craig Montell), knock-in *Gr43a-LexA* and knock-in *Gr43a-Gal4* (Hubert Amrein), and from Bloomington Drosophila Stock Center *Piezo-Gal4* (BDSC# 59266), Recombinase-M*ediated* Cassette Exchange (RMCE) gene-trap *Piezo-Gal4* (BDSC# 76658), *UAS-CD8GFP* (BDSC# 5137), *UAS-CD8RFP* (BDSC# 32218), *UAS-Trpa1* (BDSC# 26263), *Cha-Gal80* (BDSC# 60321), *UAS-CaLexA* (BDSC# 66542), *Nanchung-Gal4* (BDSC# 24903), *Inactive-Gal4* (BDSC# 36360), *Painless-Gal4* (BDSC# 27894), *Trp-Gal4* (BDSC# 36359), *Trpa1-Gal4* (BDSC# 36362), *Nompc-Gal4* (BDSC# 36361), *Ppk-Gal4* (BDSC# 32078), *UAS-DenMark* (BDSC# 33061 and 33062), *Drop-dead KO* (BDSC# 24901), *w*^*1118*^ (BDSC# 3605), *Escargot-Gal4* (BDSC# 26816), *GMR51F12-Gal4* (BDSC# 58685). *Cha-Gal80, Elav-Gal80*, and *UAS-Shibire*^*ts*^ were provided by Dragana Rogulja as published (Kitamoto, 2001; Sakai et al., 2009; Yang et al., 2009).

### Feeding analysis

Acute feeding assays were performed as previously described with modifications (Albin et al., 2015; Min et al., 2016). Twelve adult female flies were collected upon eclosion and housed in a vial with for five to seven days. Prior to testing, baseline hunger was synchronized by starving flies for 15-18 hrs in a vial containing only on a dampened kimwipe section. Regular fly food was dyed with green food coloring (McCormick, 70 µl dye per vial) and dried (24 hours). For testing, starved flies were transferred to vials containing dyed food for 30 minutes. Trials were ended by cooling the vials on ice, a feeding index was scored as described below (see quantification). For thermogenetic experiments, flies expressing Trpa1 or Shibire were maintained and starved at 18°C prior to testing. Ten minutes prior to testing, starved flies and dye-labeled food were pre-warmed to 30°C or 32°C for experiments with either Trpa1 or Shibire, and then tested as above. Feeding behavior was scored by visual inspection of ingested dye with scores given from 0 to 5 based on dye intensity, as reported previously. A feeding index was expressed by averaging the feeding scores for all flies per vial (∼12 flies). For automated analysis of feeding patterns, fasted male flies (3-5 days old) were individually introduced into chambers connected to an Expresso machine (http://public.iorodeo.com/docs/expresso/hardware_design_files.html) and feeding bouts were analyzed using Expression acquisition software (http://public.iorodeo.com/docs/expresso/device_software.html).

### Chronic studies of body weight, intestinal transit, and fecal rate, and life span

Chronic studies were performed on 5-7 day old male and female flies fed ad libitum with regular fly food. Flies were anesthetized (ice, 10 minutes) and weighed in groups of three in a 1.5 ml Eppendorf tube, with body weight expressed as the average weight per group of three. Lifespan was analyzed for a group of 12 flies by counting the number of surviving flies each day. Fecal rates were measured after feeding flies dye-colored food (dye-colored food is described above) for 1 hour, with visual inspection of abdominal dye to ensure ingestion. Flies were transferred to an empty vial containing a 1 cm X 1 cm filter paper floor for 30 minutes, and dye-labeled fecal spots on the filter paper were counted. For analysis of fecal deposition, individual data points reflect the mean behavior of ten flies. Intestinal transit was measured in flies given brief access to dye-colored food, with dye location in the intestine determined visually. A transit index was calculated based on the leading dye edge position, with scores of 1, 2, and 3 referring to dye edge in the crop/anterior midgut, middle midgut, and hindgut/anus respectively.

### Immunohistochemistry

Wholemount preparations of the gastrointestinal tract were fixed (4% paraformaldehyde, PBS, 20 min, RT), washed (2 × 5 min, PBS with 0.5% Triton X-100), permeabilized (10 min, PBS with 0.5% Triton X-100), blocked [1 hr, RT, blocking solution: 5% normal goat serum (Jackson ImmunoResearch, 005-000-121), PBS with 0.1% Triton X-100], incubated with primary antibody (1:200, blocking solution, 4°C, overnight), washed (3 × 10 min, RT, PBS with 0.1% Triton X-100), incubated with secondary antibody (1:200, PBS with 0.1% Triton X-100, 2 hr, RT), washed (3 × 10 min, RT, PBS with 0.1% Triton X-100 then 2 × 5 min, RT, PBS, mounted on a slide glass with Fluoromount-G mounting medium (Southern Biotech, 0100-01), covered with a thin coverslip, sealed with nail polish, and analyzed by confocal microscopy (Leica SP5). Primary antibodies were anti-GFP (Aves labs, Chicken, GFP-1020), anti-RFP (Rockland, Rabbit, 600-401-379), anti-Elav (Developmental Studies Hydridoma Bank, Mouse, Elav-9F8A9), and anti-Akh (Kerafast, Rabbit, EGA261). Secondary antibodies were anti-Chicken-Alexa fluor-488 (Jackson ImmunoResearch, 703-545-155), anti-Rabbit-Alexa fluor-488 (Jackson ImmunoResearch, 711-545-152), anti-Rabbit-Cy3 (Jackson ImmunoResearch, 711-165-152), anti-Rabbit-Alexa fluor-647 (Jackson ImmunoResearch, 711-605-152), anti-Mouse-Alexa fluor-647 (Jackson ImmunoResearch, 715-605-150), anti-Mouse-Alexa fuor-488 (Jackson ImmunoResearch, 715-545-150). For staining of visceral muscle and nuclei, anti-Phallodin-FITC (Sigma, P5282-1MG), anti-Phalloidin-TRITC (Sigma, P1951-1MG) and TO-PRO-3 (ThermoFisher, T3605) were added together with the secondary antibody.

### Quantification of crop size and composition

After the feeding assay, flies were fixed (4% paraformaldehyde, PBS, RT, 1 hr) and decapitated. The anterior gastrointestinal tract was surgically removed after gentle displacement of appendages and thoracic muscles. Dissected tissue was washed (3x PBS, RT, 5 min) and mounted for bright field microscopy using the “Analyze-Measure” tool in Fiji to calculate crop area. Crop muscle and cell density were quantified as detailed below. For quantification of crop muscle density, the intensity of the Phalloidin-labeled muscle fibers in ROI was divided by the total ROI area. For cell density, the number of nuclei labeled with TO-PRO-3 and counted using “Analyse-3D objects counter” function in Fiji (https://imagej.net/Fiji) was divided by total ROI area.

### Analyzing neuronal responses with CaLexA

CaLexA responses were measured in *Piezo-Gal4* or *Gr43a-Gal4* flies containing *UAS-CaLexA (LexA-VP16-NFAT, LexAop-rCD2-GFP and LexAop-CD8-GFP-2A-CD8GFP)*, and *UAS-CD8RFP*. Responses of Piezo knockout neurons were measured by introducing Piezo knockout alleles into *Piezo-Gal4*; *UAS-CaLexA; UAS-CD8RFP* flies. For sucrose and sucralose responses, flies we fed *ad libitum* with regular food, transferred to vials containing a kimwipe soaked with 10% sucrose solution or 1% sucralose solution containing green food coloring for 24 hours, and analyzed for crop distension and CaLexA expression. For water responses, flies were deprived of food and water for 6 hours, and transferred to vials containing a water-soaked kimwipe. Some flies were harvested after 15 minutes for analysis of crop distension and others were harvested after 18 hours for analysis of CaLexA expression. Control flies were placed in a vial containing a water-soaked kimwipe but no food for 24 hours and harvested for analysis. For TRPA1-mediated neuron stimulation, WT and *Piezo KO* flies bearing a *Piezo-Gal4, UAS-CaLexA (LexA-VP16-NFAT, LexAop-rCD2-GFP and LexAop-CD8-GFP-2A-CD8GFP)*, and *UAS-Trpa1* were placed in a 30°C incubator for 24 hours prior to analysis. For analysis of CaLexA expression, flies were anesthetized (ice, 10 minutes), and the anterior gastrointestinal tract was surgically removed. Dissected tissue was fixed (4% paraformaldehyde, PBS, 20 min, RT), washed (3 × 5 min, PBS), and slide mounted with Fluoromount-G mounting medium and a coverslip. Native GFP (derived from CaLexA activation) and RFP (constitutive from a Gal4-dependent reporter) fluorescence were analyzed by confocal microscopy (Leica SP5).

For quantification of CaLexA-dependent reporter in Figure 3B and S3B, intensity of GFP and RFP fluorescence was calculated per neuron and a CaLexA index expressed as GFP fluorescence divided by RFP fluorescence. For quantification of CaLexA-dependent reporter in Figure S3A and S3D, which involved flies lacking an RFP allele for neuron identification and normalization, GFP intensity was measured in the whole hypocerebral ganglion and a background subtraction was performed involving a comparably sized region of the proventriculus lacking Gal4-positive cell bodies. For S3A and S3D, background-subtracted GFP fluorescence was divided by RFP fluorescence from a control *Piezo-Gal4; UAS-CD8RFP* fly to generate a CaLexA index.

### Statistical analysis

Sample sizes from left to right (except 3B is top to bottom): Figure 2B (16, 11, 13, 10, 10 and 13, 9, 11, 9, 9), Figure 2C (19, 20, 14, 10, 13 and 12, 6, 11, 10, 12), Figure 2E (15, 20, 14, 15), Figure 3B (59, 64, 43, 67 for Gr43a neuron responses; 61, 61, 59, 66 for Piezo neuron responses; 60, 60, 33, 37 for *Piezo*-null Piezo neuron responses), Figure 4C (17, 17 and 14, 14), Figure 4D (32, 34, 33, 31), Figure 4E (21, 22 for each graph), Figure 4F (29, 22, 30, 20, 22 and 13, 13, 13, 18, 16), Figure S1B (10, 10, 10, 7), Figure S2C (8, 14, 9, 8), Figure S3A (54, 82, 44), Figure S3B (10, 10, 18, 11, 12), Figure S3C (17, 19, 28), Figure S3D (22, 23), Figure S4B (11, 11, 10, 11, 11, 10), Figure S4C (13, 16, 17), Figure S4D (8, 7, 7 for WT, Drd KO and Piezo KO), Figure S4E (9, 10, 10), Figure S4F (7, 10), Figure S4G (7, 10), Figure S4H (15, 9), Figure S4I (14, 15, 15, 15, 20 and 12, 12, 12, 12, 12), Figure S4J (10, 9, 9, 9, 9, 9 and 17, 15, 14, 16, 16, 10). Data in graphs are represented as means ± SEM. Statistical significance was analyzed by ANOVA Dunnett’s multiple comparison test or unpaired t-test using Prism 8 software (Graphpad), as indicated in figure legends.

